# ‘Big Isotopic Data’ link millet consumption under Lombard rule in early medieval northern Italy

**DOI:** 10.1101/2025.08.28.672877

**Authors:** Carlo Cocozza

**Affiliations:** Domestication and Anthropogenic Evolution Research Group (DAE), Max Planck Institute of Geoanthropology, Kahlaische Str. 10, Jena, 07745, Germany; ArchaeoBioCenter (ABC), Ludwig-Maximilians-Universität München, Geschwister-Scholl-Platz 1, München, 80539, Germany

**Keywords:** Stable isotopes, Millet, Bayesian Modelling, Early Medieval Italy, Lombards

## Abstract

Archaeobotanical and biomolecular evidence from northern Italy has highlighted a significant increase in millet cultivation during the early medieval period. This study applies Bayesian spatiotemporal modelling to the Isotòpia and CIMA databases, which collected isotopic measurements on bioarchaeological remains from ancient and medieval Europe, respectively, to investigate a debated historical link between this agricultural shift and the Lombard rule in northern Italy (568-774 CE). Results confirm that this period coincides with an increased reliance on millet, particularly in regions associated with the Po Valley. This shift is primarily attributed to the need for food security in a politically fragmented and unstable environment, with millet offering reliable yields due to its phenotypic traits for tolerance. While political agenda and environmental factors appear to impact millet cultivation to varying degrees, specific links with social or ethnic affiliations remain largely unclear and require additional research. The approach used in this study demonstrates the potential for ‘Big Isotopic Data’ to identify major historical trends.

## Introduction

Millet is a collective term used to describe small-seeded plants that share similar phenotypic characteristics, such as being drought-resistant, fast-growing, and capable of thriving in poor soils (Food and Agriculture Organization of the United Nations (FAO), 1995; James et al., 2011; Kirleis et al., 2022). Millet also provides high yields that can be stored for extended periods, making it a reliable food source when other crops fail (Food and Agriculture Organization of the United Nations (FAO), 1995; James et al., 2011; Kirleis et al., 2022). Highly mobile communities have historically chosen to cultivate this crop, given the need of fast-growing and durable food sources (Food and Agriculture Organization of the United Nations (FAO), 1995; James et al., 2011; Kirleis et al., 2022). With its relatively high nutritional content, limited economic effort, and low carbon and water footprints, scientists and international organizations advocate for the cultivation of millet to ensure food security in regions facing climatic and environmental challenges (Antony Ceasar & Maharajan, 2022; Food and Agriculture Organization of the United Nations (FAO), 1995; Saleh et al., 2013; Samtani et al., 2024). However, despite the nutritional, economic, and sustainability benefits offered by this crop, millet cultivation remains largely concentrated in specific regions of the Global South, suggesting that cultural and historical dynamics may impact its distribution (Deevi et al., 2024).

Archaeological research has explored domestication, cultivation, distribution, and consumption of millet among ancient populations (Hermes et al., 2019; Miller et al., 2016; Motuzaitė Matuzevičiūtė & Laužikas, 2023; Orfanou et al., 2024; Wilkin et al., 2020). Yet, significant historical gaps remain in the understanding of how the interplay of environmental conditions, cultural preferences, socioeconomic structures, and political agendas influenced millet agricultural practices among ancient human groups. The Early Middle Ages (circa 500-1000 CE) in northern Italy provide an ideal context for exploring these dynamics, as the region experienced significant cultural, socio-economic, and political transformations following the fall of the Western Roman Empire in 476 CE (Ward-Perkins, 2006; Wickham, 2006, 2010). While broomcorn (*Panicum miliaceum*) and foxtail (*Setaria Italica*) millet were certainly known in Roman Italy, previous studies have shown that these crops were neither a staple in the Roman diet nor central to their agricultural economy (Bosi et al., 2020; Heinrich et al., 2021; Murphy, 2016; Spurr, 1983). Conversely, earlier research has demonstrated that millet gained prominence, specifically in northern Italy, during the early medieval period (Bosi et al., 2020; Castiglioni & Rottoli, 2013; Ganzarolli et al., 2018; Iacumin et al., 2014; Montanari, 1988). Political upheavals following the alternate and fragmented rule of Germanic groups, such as the Heruli, Ostrogoths, Lombards, and Franks, and that of the Eastern Roman (also known as Byzantine) Empire, have reshaped the socioeconomic and cultural landscape of the region. A pivotal event has certainly been the Lombard migration and establishment of a kingdom in 568 CE, after a century marked by political instability, wars, the first plague pandemic, and famines (Gasparri, 2016; Jacobsen, 2009; Jarnut, 1982; Pohl, 2008a, 2008b). While also controlling strategic regions in central and southern Italy (*Langobardia Minor*) through de facto autonomous duchies, the Lombard rule was primarily focused in northern Italy (*Langobardia Maior*) (Gasparri, 2016; Jarnut, 1982). Whether the Lombard migration from central Europe, where millet consumption among Germanic and Eurasian populations is known (Alt et al., 2014; Faragó et al., 2022; Gyulai, 2014; Hakenbeck et al., 2017; Plecerová et al., 2020), directly impacted the cultivation of this crop in northern Italy remains a topic of debate (Bosi et al., 2020; Castiglioni & Rottoli, 2013; Ganzarolli et al., 2018; Iacumin et al., 2014; Marinato, 2019; Montanari, 1988; Riccomi et al., 2020).

Human-millet relationship in ancient populations has been documented through written records, archaeobotanical studies of charred remains, and biomolecular identification of miliacin compound markers in cooking vessels (Castiglioni & Rottoli, 2013; Ganzarolli et al., 2018; Gyulai, 2014; Miller et al., 2016; Spurr, 1983; Taché et al., 2021). Additionally, stable carbon (δ^13^C) isotope ratios measured in human remains offer direct insights on consumption of millet or millet-fed animals (Cerling et al., 1997; Lee-Thorp, 2008). Millet, a C_4_ group of plants, exhibits distinctly higher δ^13^C values than C_3_ plants, including most common cereals (such as wheat, barley, spelt, rye, and rice), pulses, vegetables, and fruit, in addition to terrestrial animals thriving in a C_3_ environment (Cerling et al., 1997; Lee-Thorp, 2008). An exception is marine resources, which typically exhibit comparatively high δ^13^C values that overlap with millet stable carbon isotopic ratios. Whereas freshwater resources are often ^13^C-depleted, hence presenting relatively low δ^13^C values typically overlapping or being more negative than C_3_ terrestrial foods (Lee-Thorp, 2008; Mion et al., 2022). To enhance dietary reconstruction and account for potential aquatic food consumption, researchers often analyse stable carbon and nitrogen (δ^15^N) isotope ratios together^32^. Stable nitrogen isotopes are a proxy reflecting the full protein segment of a diet. It typically shows comparatively higher values in omnivores (e.g. pigs) and aquatic animals than in herbivores or plants, in addition to informing on agricultural management practices (e.g. manuring) and environmental conditions (e.g. aridity) (Bogaard et al., 2007; Lee-Thorp, 2008; Styring et al., 2016; Szpak, 2014). This combined approach has been widely used to build individual dietary osteobiographies and reconstruct subsistence practices in early medieval communities (Alt et al., 2014; Faragó et al., 2022; Hakenbeck et al., 2017; Iacumin et al., 2014; Plecerová et al., 2020; Riccomi et al., 2020).

Recently, collaborative efforts have aggregated large volumes of isotopic information from various regions and time periods, enhancing research potential and documenting significant dietary shifts across different spatiotemporal scales (Billings et al., 2025; Cocozza et al., 2022; Ebert et al., 2024; Farese et al., 2023; Formichella et al., 2024; Goldstein et al., 2022; Mantile et al., 2023; Pezo-Lanfranco et al., 2024). In this study, spatiotemporal Bayesian modelling was applied to published isotopic data compiled within the Isotòpia (Formichella et al., 2023, 2024) and CIMA (Cocozza, Cirelli, et al., 2021; Cocozza et al., 2022) databases, which focus on ancient and medieval Europe and the Mediterranean, respectively. The aim was that of exploring historical links between the Lombard migration to northern Italy and the documented increase in millet consumption in the region during the Early Middle Ages.

## Materials and Methods

The Isotòpia (Formichella et al., 2023, 2024) and CIMA (Cocozza, Cirelli, et al., 2021; Cocozza et al., 2022) databases collectively compile published isotopic data originating from Europe and the Mediterranean, covering a combined chronological span from 800 BCE to 1500 CE. Both databases are hosted on the Pandora data platform (https://pandoradata.earth/) and are part of the IsoMemo network (https://isomemo.com/), a collaborative open-access initiative of independent isotopic databases. Isotòpia is accessible via https://www.doi.org/10.48493/m0m0-b436, while CIMA can be found at https://www.doi.org/10.48493/s9nf-1q80.

The Pandora & IsoMemo initiative developed several open-access, user-friendly R-based modelling tools tailored for historical data analysis (Cocozza et al., 2023, 2025; Cocozza, Fernandes, et al., 2021; Cubas et al., 2020; Wilkin et al., 2020). To explore spatiotemporal variations in isotopic data for early medieval Italy and central Europe, δ^13^C_collagen_, δ^13^C_bioapatite_, and δ^15^N_collagen_ isotopic records of human individuals registered in Isotòpia and CIMA were Bayesian modelled through available options provided by the Data Search and Spatiotemporal Modelling (DSSM) application. The code for DSSM is accessible on GitHub at https://github.com/Pandora-IsoMemo/DSSM. The TimeR (Cocozza et al., 2022) modelling tool available in DSSM (v. 25.1.0.1) was employed to identify temporal human isotopic variations across central Europe and northern Italy during the late Roman and early medieval periods. TimeR is a Bayesian geostatistical model, that offers a smoothing estimate for a “dependent” variable across both time and space (Cocozza et al., 2022). Additionally, the KernelTimeR (Cocozza et al., 2022) modelling tool (v. 25.1.0.1) was employed to group northern Italian sites based on their spatial distribution.

To account for metabolic routing differences (Fernandes et al., 2012; Lee-Thorp, 2008), direct comparison between human δ^13^C_collagen_ and δ^13^C_bioapatite_ values was provided through the conversion of the latter into the former using the mathematical formula proposed by Lee-Thorp et al., (1989) for omnivores. To take into account additional uncertainties introduced by this conversion, a conservative uncertainty margin of 1‰ was considered for the modelling of converted values. To ensure the integrity of the modelling results, data that might have been compromised by diagenesis or poor measurement quality was selectively excluded. Records that did not meet the atomic C:N ratio criteria of 2.9-3.6 (DeNiro, 1985), as well as those with %C and %N elemental composition below 13% and 4.8%, respectively, were excluded (Ambrose, 1990). Moreover, collagen data whose %C and %N elemental composition was above 50% and 20%, respectively, as well as those presenting δ^13^C values below -23‰ and/or δ^15^N values below 5‰ were removed since they are likely to reflect collagen contamination or instrumental/calibration errors (Billings et al., 2025). Infant records were also excluded given that their diet may potentially differ from the one practiced by the adult population. For the same reason, isotopic data from tooth samples and petrous bones of adults were also filtered out, given that these reflect childhood dietary intake (Fuller et al., 2003; Jørkov et al., 2009). A filter was applied to include only records whose chronological range overlapped with the first millennium CE. Finally, in cases of data duplication between the Isotòpia and CIMA databases, entries from the former were preferentially excluded to avoid redundancy.

## Results and Discussion

Figure 1 illustrates spatiotemporal trends for modelled adult human bone collagen stable carbon (**Figure 1a**) and nitrogen (**Figure 1b**) isotopic data across northern Italy and central Europe from 450 to 850 CE, displayed at 50-year intervals. In a broad area roughly corresponding to continental parts of modern-day Croatia, Serbia, Hungary, and Romania, relatively high δ^13^C and δ^15^N values are consistently observed throughout the entire period of analysis, with little temporal variations. The only exception is an increase in δ^13^C values after 700 CE. Such isotopic values may indicate a diet either rich in marine resources or composed of a mix of ^15^N-enriched millet-fed terrestrial animals paired with significant millet consumption. Given the lack of evidence for marine fish consumption in this landlocked area during the early medieval period, it is inferred that high δ^13^C values are primarily linked to millet, a crop present in the region since the late Bronze Age (Kirleis et al., 2022; Miller et al., 2016; Orfanou et al., 2024). In northern Italy, while δ^15^N values show almost no variation, δ^13^C values exhibit significant temporal shifts across much of the region. From approximately 550 to 700 CE, δ^13^C values rise, stabilise around 750 CE, and gradually decline by 800 to 850 CE. Unlike in central Europe, defining a dietary mixing line in northern Italy is more challenging, due to the likely presence of marine resources. Mediterranean fish, while displaying considerable variability in δ^15^N values, present on average lower stable nitrogen ratios values compared to their Atlantic counterparts (Mion et al., 2022). This variability complicates precise dietary assessments based solely on bulk stable isotope analysis (Soncin et al., 2021). Nevertheless, archaeobotanical and biomolecular evidence from the Early Middle Ages confidently associates the significant rise in δ^13^C with shifts in millet or millet-fed animal consumption (Bosi et al., 2020; Castiglioni & Rottoli, 2013; Ganzarolli et al., 2018; Iacumin et al., 2014; Marinato, 2019).

Results from central Europe offer valuable insights into the relationship between different populations and millet consumption. While frequent migrations and political upheavals among Eurasian and Germanic populations complicate ethnic attribution of isotopic variations, it is observed a clear east-west divide in δ^13^C values from 5^th^-6^th^ centuries Hungary that aligns with both environmental and political borders. In the eastern part, the Great Hungarian Plains were encompassed into the Hunnic confederation until its collapse in the mid-5^th^ century, after which the Germanic Gepids, former Hunnic subjects, established their kingdom (Heather, 1995). Here comparatively high δ^13^C and δ^15^N values suggest that millet was a dietary staple. It is possible that millet was ^15^N-enriched due to arid conditions in the region, and that its consumption among humans was coupled with livestock products from millet-fed sheep, goats, and cattle (Hakenbeck et al., 2017). This dietary pattern aligns with the pastoralist economy typical of the Huns, while the Gepids may have little changed the local agricultural organisation (Heather, 1995). The region west of the Danube (Transdanubia) transitioned from the Roman province of Pannonia to short-lived Hunnic control and later hosted various Germanic tribes (Heather, 1995). This includes the Lombards, who moved from southern Moravia in the first half of the 6^th^ century, as documented historically and confirmed by archaeological evidence (Jarnut, 1982; Pohl, 2008b; Šmerda, 2016; Vida, 2008). Isotopic results from the region indicate a mixed diet of C_3_ plants and millet, along with ^15^N-enriched animal sources. The moderate presence of millet in Lombard diets might have originated before their migration to Transdanubia, as isotopic data from Lombard sites in Kyjov (Czech Republic) (Plecerová et al., 2020) and Szólád (Hungary) (Alt et al., 2014) show no significant differences. In the second half of the 6^th^ century, the Lombards defeated the Gepids and ceded control of central Europe to the Avars, a Eurasian population (Gnecchi-Ruscone et al., 2022; Pohl, 2018), while they themselves migrated to Italy to establish their own kingdom (Gasparri, 2016; Jarnut, 1982; Pohl, 2008a, 2008b). Bayesian-modelled isotopic results align with this historical transition, showing an increase and spread in δ^13^C values that correspond with the Avars’ expansion further into central Europe up to the Carinthian Alps until the 9^th^ century (Pohl, 2018). This indicates that millet kept and even increased its central role in the farming economy of the region, possibly due to cultural links with populations residing in the Avar Khaganate, and likely as part of a defined political agenda.

**Figure 1.**
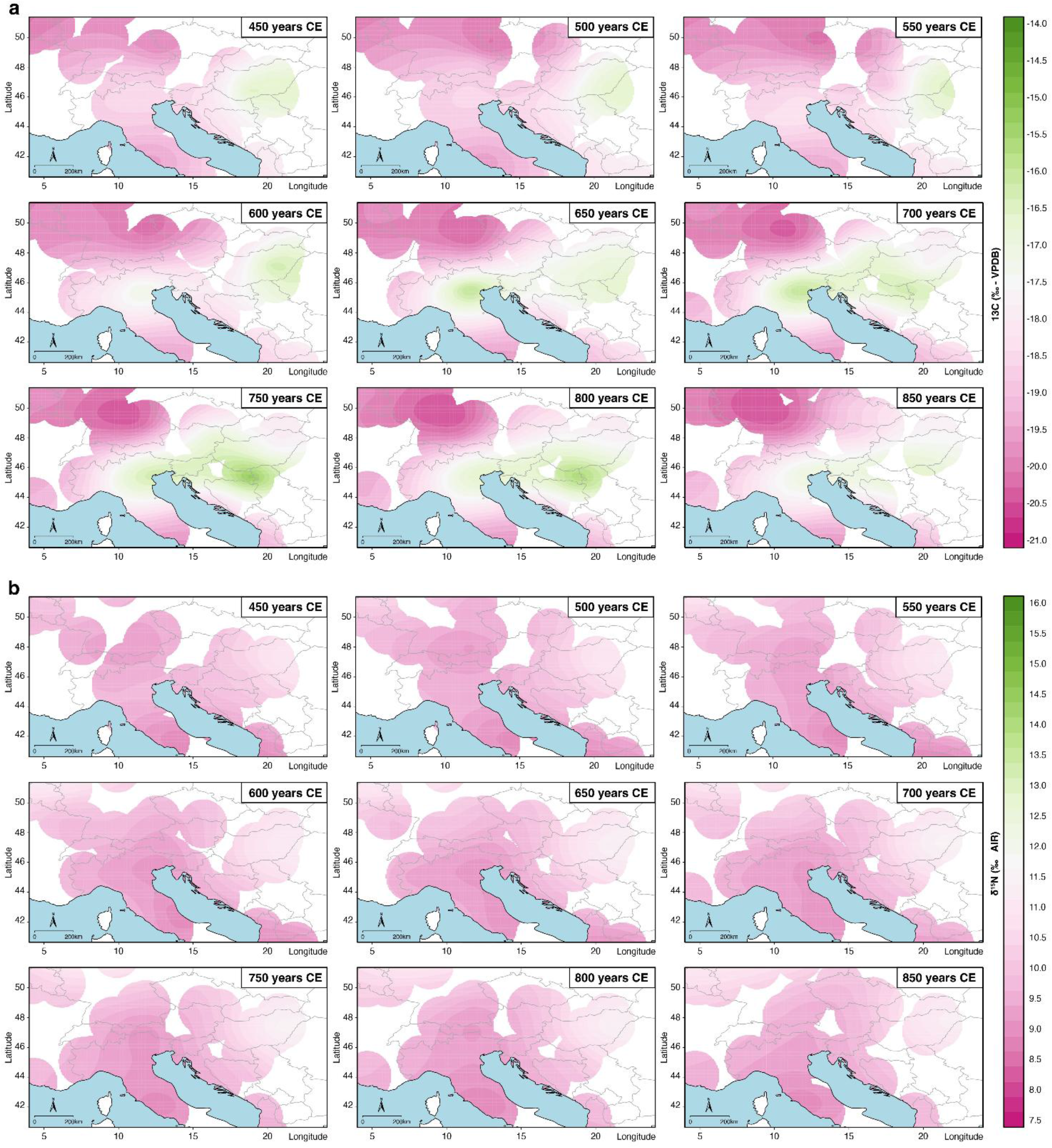
Bayesian spatiotemporal δ^13^C (**a**) and δ^15^N (**b**) models for adult human bone collagen across northern Italy and central Europe from 450 to 850 CE, presented at 50-year intervals. Maps generated via the TimeR modelling option (v. 25.1.0.1) available in Data Search and Spatiotemporal Modelling application (https://github.com/Pandora-IsoMemo/DSSM).

In 568 CE the Lombards established a kingdom in Italy that encompassed most of the northern region (*Langobardia Maior*) along with strategic areas in central and southern parts (*Langobardia Minor*), such as the Duchies of Spoleto and Benevento (Gasparri, 2016; Jarnut, 1982). By 774 CE, Charlemagne absorbed the Lombard kingdom into the Frankish realm (Gasparri, 2016; Jarnut, 1982). The isotopic shifts observed in the Bayesian modelled results chronologically align with the Lombard period of rule. While δ^13^C values from 450 to 550 CE also possibly indicate moderate millet consumption, this becomes increasingly significant after 600 CE and potentially a staple in the diet of northern Italian populations. The stable carbon isotope values rise substantially until around 700/750 CE, before declining by 800 and 850 CE, suggesting that millet lost its predominance in the farming economy of the region during the Frankish domination.

A visual examination of data in northern Italy highlighted a distinct concentration of sites in the northeast, while those in the northwest appeared more sporadic. To formally identify spatial clusters, a k-means clustering was employed via the KernelTimeR (Cocozza et al., 2022) tool. To do so, the chronological range of each Italian archaeological site north of modern Umbria, Marche, and Lazio was considered (**Figure 2a-b**). The spatial centroid coordinates of these clusters were then compared using TimeR (Cocozza et al., 2022) to evaluate temporal isotopic shifts between cluster areas (**Figure 2c-d**). This analysis robustly illustrates the Lombards’ impact on the millet agricultural economy in northern Italy. A clearly positive δ^13^C trend is evident for Cluster East, prominently aligning with the Lombard ruling period, whereas δ^13^C values for Cluster West show a slight increase, beginning alongside Cluster East and extending for a longer duration

**Figure 2.**
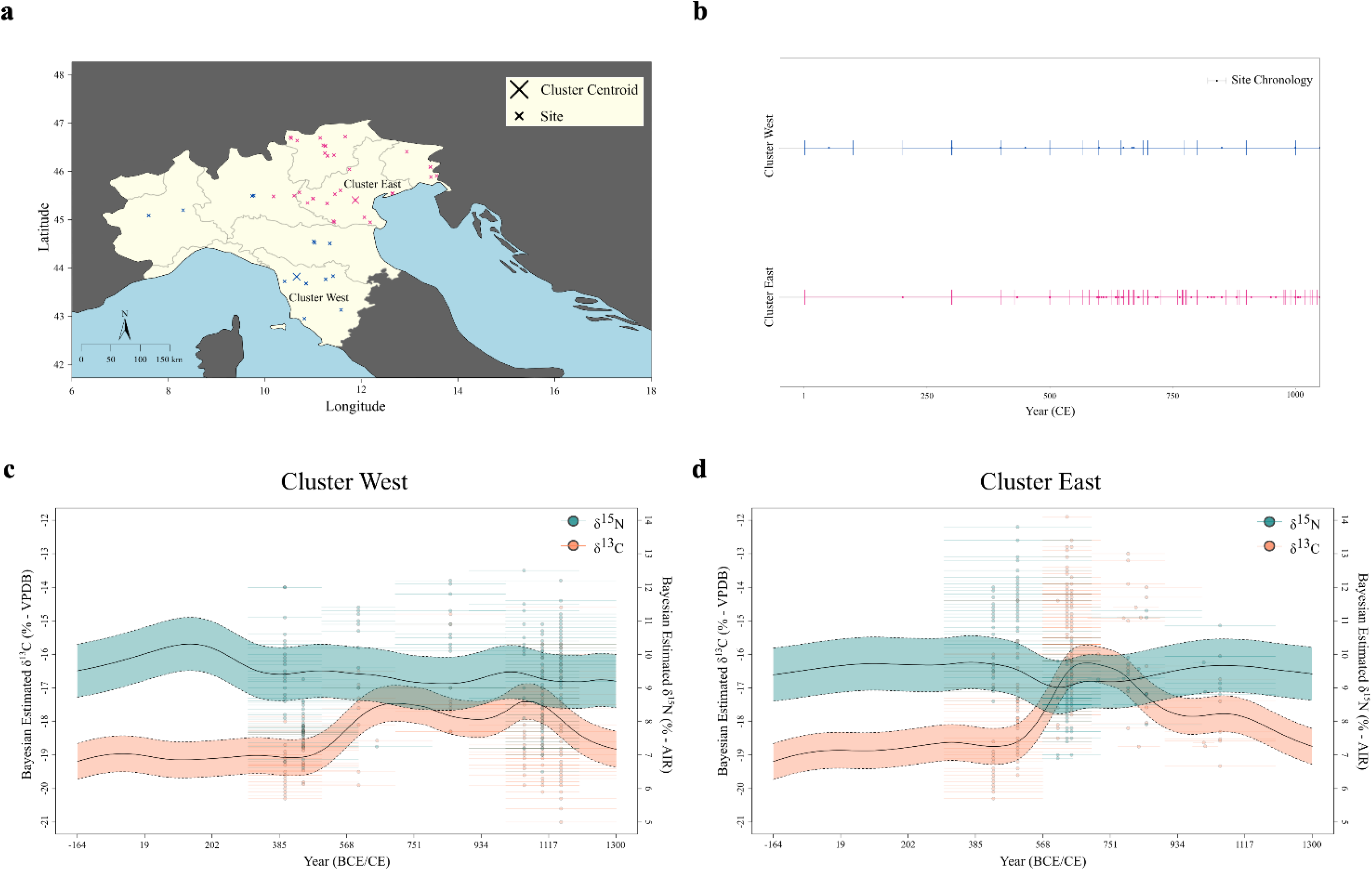
**a)** Spatial cluster analysis of Early Medieval sites from northern Italy; **b**) Temporal distribution of these sites across the identified clusters; **c**) Temporal variation of δ^13^C and δ^15^N isotopic values for Cluster West; **d**) Temporal variation of δ^13^C and δ^15^N isotopic values for Cluster East. Figure **a** and **b** were produced via the KernelTimeR modelling option (v. 25.1.0.1) available in Data Search and Spatiotemporal Modelling application (https://github.com/Pandora-IsoMemo/DSSM). While Figure **c** and **d** were generated via the TimeR tool (v. 25.1.0.1).

Early medieval Italy was characterised by significant political fragmentation (Gasparri, 2016; Jarnut, 1982). In the first decades of the Lombard kingdom, regions such as the northern coastal areas, the region surrounding the former Western Roman capital of Ravenna, a land corridor connecting Ravenna to Rome, Latium, and significant portions of southern Italy remained under the control of the Eastern Roman Empire. Although the Lombard domains gradually expanded during the 7^th^ and early 8^th^ centuries, ongoing warfare and internal political upheavals fostered socioeconomic and political instability. This possibly accentuated the agricultural regionalisation process that Italy was already experiencing since the fall of the Western Roman Empire (MacKinnon, 2019; Montanari, 1988; Riccomi et al., 2020). Favourable climatic conditions and relative political stability supported the traditional prominent cultivation of C_3_ cereals like wheat and barley in southern and central regions, including sites from the *Langobardia Minor*, as supported by isotopic evidence (Bernardini et al., 2021; Nitsch, 2012; Tafuri et al., 2018). Whereas northern Italy largely transitioned to millet production (Montanari, 1988). The motivation behind this shift remains unclear; however, it could have been promoted – whether actively or not – by the Lombard elites, who were familiar with millet from their past in central Europe. On the other hand, a bottom-up initiative may have also been possible, given an agricultural system that granted local landowners substantial autonomy. Lacking central authority support, millet could have become a pivotal crop offering food security in a context of financial uncertainty and political instability. In addition, harsher climatic conditions than those from other Italian regions and environmental-specific features also possibly played a role in crop selection (Riccomi et al., 2020). An increase in δ^13^C values during Lombard rule, as demonstrated by Bayesian spatiotemporal modelling, is notably associated with the Po Valley, a vast alluvial plain encompassing much of present-day Friuli Venezia Giulia, Veneto, Emilia-Romagna, Lombardy, and parts of Piedmont (**Figure 1a** and **2d**). Conversely, results from the hilly terrains of Tuscany, the central Apennines, and toward the Alps indicate a more moderate increase in δ^13^C values, suggesting that millet was accompanied by other C_3_ crops (**Figure 1a** and **2c**).

Figure 3 illustrates the distribution of individual adult human bone collagen δ^13^C and δ^15^N values by cultural affiliation in northern Italy. The data used in this plot is the same employed for Bayesian spatiotemporal modelling that generated **Figure 1** and only pertain to northern Italian sites as for **Figure 2**. The “Roman” label is used to indicate northern Italian populations that are clearly identified as the local ‘Roman’ substratum in original publications, albeit these may include individuals predating 568 CE (Maxwell, 2019; Temkina, 2021). Whereas the “Lombard” label refers to populations that were identified as culturally Lombard (Amorim et al., 2018; Fiorin et al., 2021; Iacumin et al., 2014; Laffranchi et al., 2020; Marinato, 2014, 2017, 2018; Maxwell, 2019; Paladin et al., 2020; Ricci et al., 2012; Riccomi et al., 2020; Tian et al., 2024), although it may include ethnically Roman individuals buried in Lombard site. The spatial cluster division is here highlighted only for Lombards, whilst all Romans belong to sites affiliated to Cluster East (**Figure 2**).

**Figure 3.**
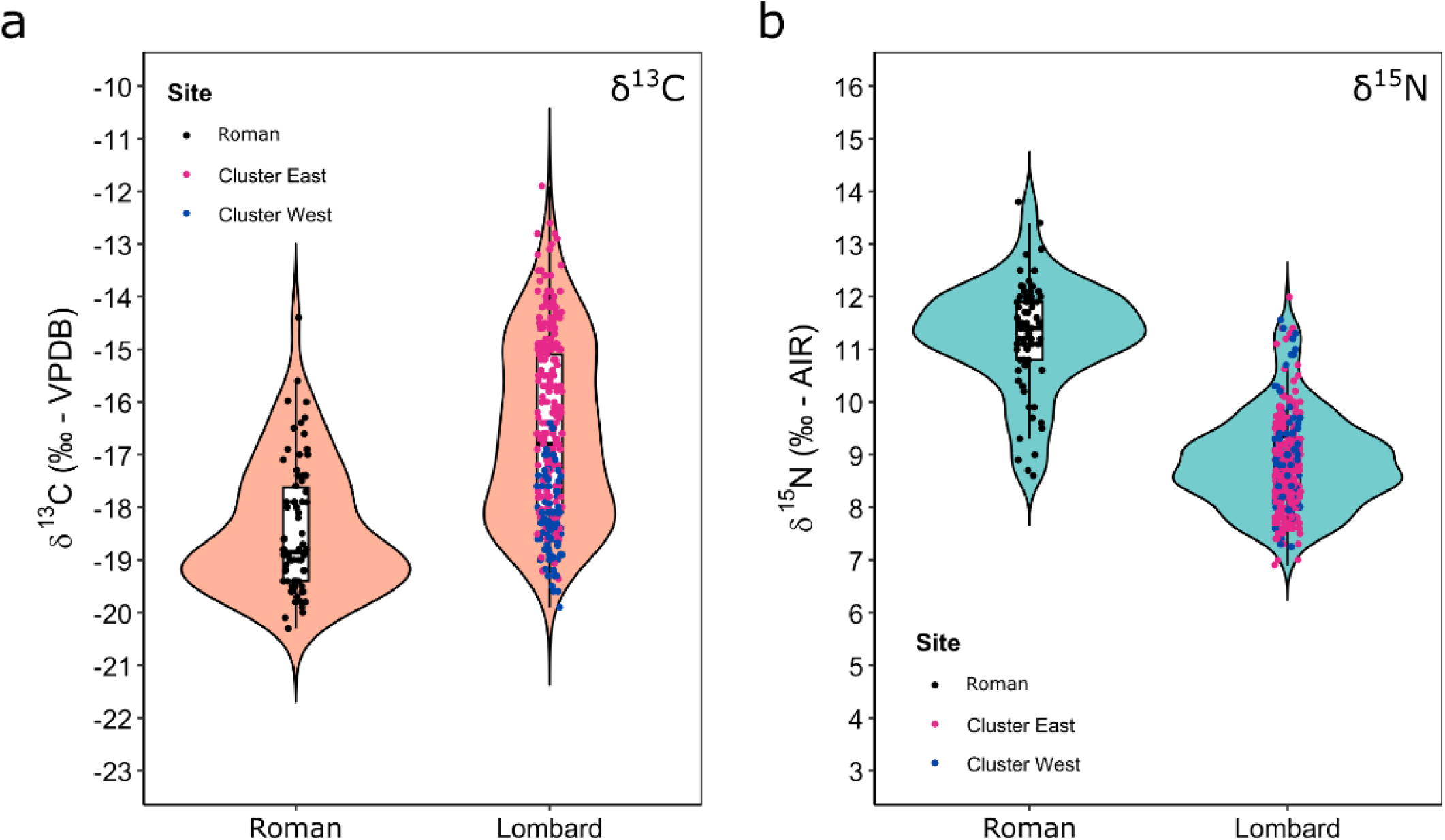
Violin plots showing the distribution of adult human bone collagen δ^13^C (**a**) and δ^15^N (**b**) values for culturally Roman or Lombard individuals.

The violin plots in **Figure 3** reveal significant variability in δ^13^C and δ^15^N values among culturally Roman and Lombard populations in northern Italy. The comparison indicates that Lombard δ^13^C values exhibit a broader distribution than those of the Romans. This variability may be attributed to differing levels of millet consumption, possibly linked to environmental variations discussed previously or distinct dietary practices across social classes. Differences in status may also be based on ethnicity, especially during the early centuries of Lombard rule, which marked a significant social divide between Romans and Lombards (Gasparri, 2016). The distinct separation into Cluster East and West in Lombard δ^13^C values aligns with trends observed in Bayesian models for most individuals and may enhance a predominantly environmental explanation. The narrower range of δ^15^N values among the Lombards, which does not offer distinct cluster patterns, supports the hypothesis that variations in δ^13^C values are likely due to differing dietary proportions of millet rather than the presence of marine resources. Roman δ^13^C values remain broadly distributed, suggesting the inclusion of millet in their diet and hinting at a widespread agricultural practice that went beyond the borders of the Lombard kingdom. Nonetheless, the mirroring distribution of δ^15^N values may imply that δ^13^C values may also include varying proportions of marine resource consumption at intra- or inter-site levels, possibly associated with social status.

## Conclusion

The ‘Big Isotopic Data’ approach employed in this study revealed significant links between isotopic variations and historical developments following the Lombard migration to northern Italy during the Early Middle Ages. While site-specific differences likely existed, a general trend indicates that a high consumption of millet was concurrent with the movements of Eurasian and Germanic populations since the 5^th^ century (or earlier) in central Europe. In the aftermath of the Western Roman Empire’s collapse, these communities either adopted or were compelled to adopt agricultural practices that were more suitable to their lifestyles and local environmental conditions. The agrarian system of the Roman Empire, which had heavily relied on wheat and barley, was no longer sustainable within the prevailing political instability, with few exceptions. Consequently, millet production and consumption rose to prominence due to its versatility and adaptability, providing essential food security to early medieval communities. This trend reaches a peak with the Lombards in northern Italy, who may have enhanced the role of millet through a farming system that gave large autonomies to land owners who, in turn, heavily relied on this crop. However, it remains unclear whether the Lombards actively promoted millet cultivation based on their previous experiences in Central Europe or if this emerged independently as a response by the northern Italian populations to new circumstances.

Future research on early medieval Italy should prioritise improving chronological precision, identifying status affiliations (in terms of both wealth and ethnicity), and addressing existing gaps in spatiotemporal isotopic studies, particularly those involving Eastern Roman regions. This would enable in-depth intra- and inter-socioeconomic comparisons concerning dietary practices and offer answers to historical aspects that remain unclear. Moreover, to improve the accuracy of direct dietary reconstructions, additional isotopic analyses of faunal and charred plant remains are essential, as this study was unable to explore these elements in detail.

## Acknowledgements

I wish to thank Michelle O’Reilly for her valuable assistance with **Figure 1**. I also extend my gratitude to Dr. Ricardo Fernandes, Dr. Robert Spengler, Dr. Silvia Soncin, Prof. Wolf-Rüdiger Teegen, Prof. Bernd Päffgen, and Daniela Coppola for their insightful feedback during the development of this manuscript.

## References

Alt, K. W., Knipper, C., Peters, D., Müller, W., Maurer, A.-F., Kollig, I., Nicklisch, N., Müller, C., Karimnia, S., Brandt, G., Roth, C., Rosner, M., Mende, B., Schöne, B. R., Vida, T., & von Freeden, U. (2014). Lombards on the Move – An Integrative Study of the Migration Period Cemetery at Szólád, Hungary. PLoS ONE, 9(11), Article 11. 10.1371/journal.pone.0110793

Ambrose, S. H. (1990). Preparation and characterization of bone and tooth collagen for isotopic analysis. Journal of Archaeological Science, 17(4), Article 4. 10.1016/0305-4403(90)90007-R

Amorim, C. E. G., Vai, S., Posth, C., Modi, A., Koncz, I., Hakenbeck, S., La Rocca, M. C., Mende, B., Bobo, D., Pohl, W., Baricco, L. P., Bedini, E., Francalacci, P., Giostra, C., Vida, T., Winger, D., von Freeden, U., Ghirotto, S., Lari, M., … Veeramah, K. R. (2018). Understanding 6th-century barbarian social organization and migration through paleogenomics. Nature Communications, 9(1), Article 1. 10.1038/s41467-018-06024-4

Antony Ceasar, S., & Maharajan, T. (2022). The role of millets in attaining United Nation’s sustainable developmental goals. PLANTS, PEOPLE, PLANET, 4(4), 345–349. 10.1002/ppp3.10254

Bernardini, S., Asrat Mogesie, S., Micarelli, I., Manzi, G., & Tafuri, M. A. (2021). Contribution to Longobard dietary studies: Stable carbon and nitrogen isotope data from Castel Trosino (6th-8th c. CE, Ascoli Piceno, central Italy). Data in Brief, 38, 107290. 10.1016/j.dib.2021.107290

Billings, T. N., Scott, E., Cocozza, C., Hixon, S., Boivin, N., Roberts, P., Spengler, R. N., & Fernandes, R. (2025). The North American Repository for Archaeological Isotopes. Scientific Data, 12(1), 50. 10.1038/s41597-024-04175-2

Bogaard, A., Heaton, T. H. E., Poulton, P., & Merbach, I. (2007). The impact of manuring on nitrogen isotope ratios in cereals: Archaeological implications for reconstruction of diet and crop management practices. Journal of Archaeological Science, 34(3), Article 3. 10.1016/j.jas.2006.04.009

Bosi, G., Castiglioni, E., Rinaldi, R., Mazzanti, M., Marchesini, M., & Rottoli, M. (2020). Archaeobotanical evidence of food plants in Northern Italy during the Roman period. Vegetation History and Archaeobotany, 29(6), 681–697. 10.1007/s00334-020-00772-4

Castiglioni, E., & Rottoli, M. (2013). Broomcorn millet, foxtail millet and sorghum in north Italian Early Medieval sites. Post-Classical Archaeologies, 3, 131–144.

Cerling, T. E., Harris, J. M., MacFadden, B. J., Leakey, M. G., Quade, J., Eisenmann, V., & Ehleringer, J. R. (1997). Global vegetation change through the Miocene/Pliocene boundary. Nature, 389(6647), 153– 158.10.1038/38229

Cocozza, C., Cirelli, E., Groß, M., Teegen, W.-R., & Fernandes, R. (2021). Compendium Isotoporum Medii Aevi (CIMA) [Computer software]. 10.48493/s9nf-1q80

Cocozza, C., Cirelli, E., Groß, M., Teegen, W.-R., & Fernandes, R. (2022). Presenting the Compendium Isotoporum Medii Aevi, a Multi-Isotope Database for Medieval Europe. Scientific Data, 9(1), Article 1. 10.1038/s41597-022-01462-8

Cocozza, C., Fernandes, R., Ughi, A., Groß, M., & Alexander, M. M. (2021). Investigating infant feeding strategies at Roman Bainesse through Bayesian modelling of incremental dentine isotopic data. International Journal of Osteoarchaeology, 31(3), Article 3. 10.1002/oa.2962

Cocozza, C., Harris, A. J. T., Formichella, G., Pedrucci, G., Rossi, P. F., D’Alessio, A., Amoretti, V., Zuchtriegel, G., O’Reilly, M., Mantile, N., Panella, S., Tafuri, M. A., Altieri, S., di Cicco, M. R., Fernandes, R., & Lubritto, C. (2025). High-resolution isotopic data link settlement complexification to infant diets within the Roman Empire. PNAS Nexus, 4(1), pgae566. 10.1093/pnasnexus/pgae566

Cocozza, C., Teegen, W.-R., Vigliarolo, I., Favia, P., Giuliani, R., Muntoni, I. M., Oione, D., Clemens, L., Groß, M., Roberts, P., Lubritto, C., & Fernandes, R. (2023). A Bayesian multi-proxy contribution to the socioeconomic, political, and cultural history of Late Medieval Capitanata (southern Italy). Scientific Reports, 13, 4078. 10.1038/s41598-023-30706-9

Cubas, M., Lucquin, A., Robson, H. K., Colonese, A. C., Arias, P., Aubry, B., Billard, C., Jan, D., Diniz, M., Fernandes, R., Fábregas Valcarce, R., Germain-Vallée, C., Juhel, L., de Lombera-Hermida, A., Marcigny, C., Mazet, S., Marchand, G., Neves, C., Ontañón-Peredo, R., … Craig, O. E. (2020). Latitudinal gradient in dairy production with the introduction of farming in Atlantic Europe. Nature Communications, 11(1), Article 1. 10.1038/s41467-020-15907-4

Deevi, K. C., Swamikannu, N., & Padmanabhan, J. (2024). Global Millet Trends, Outlook, Challenges, and Opportunities. In R. K. Srivastava, C. T. Satyavathi, & R. K. Varshney (Eds.), The Pearl Millet Genome (pp. 1–14). Springer International Publishing. 10.1007/978-3-031-56976-0_1

DeNiro, M. J. (1985). Postmortem preservation and alteration of in vivo bone collagen isotope ratios in relation to palaeodietary reconstruction. Nature, 317(6040), 806–809. 10.1038/317806a0

Ebert, C. E., Hixon, S. W., Buckley, G. M., George, R. J., Pacheco-Fores, S. I., Palomo, J. M., Sharpe, A. E., Solís-Torres, Ó.R., Davis, J. B., Fernandes, R., & Kennett, D. J. (2024). The Caribbean and Mesoamerica Biogeochemical Isotope Overview (CAMBIO). Scientific Data, 11(1), 349. 10.1038/s41597-024-03167-6

Faragó, N., Gáll, E., Gulyás, B., Marcsik, A., Molnár, E., Bárány, A., & Szenthe, G. (2022). Dietary and cultural differences between neighbouring communities: A case study on the early medieval Carpathian Basin (Avar and post-Avar period, 7th–9th/10th centuries AD). Journal of Archaeological Science: Reports, 42, 103361. 10.1016/j.jasrep.2022.103361

Farese, M., Soncin, S., Robb, J., Fernandes, R., & Tafuri, M. A. (2023). The Mediterranean archive of isotopic data, a dataset to explore lifeways from the Neolithic to the Iron Age. Scientific Data, 10(1), 917. 10.1038/s41597-023-02783-y

Fernandes, R., Nadeau, M.-J., & Grootes, P. M. (2012). Macronutrient-based model for dietary carbon routing in bone collagen and bioapatite. Archaeological and Anthropological Sciences, 4(4), Article 4. 10.1007/s12520-012-0102-7

Fiorin, E., Moore, J., Montgomery, J., Lippi, M. M., Nowell, G., & Forlin, P. (2021). Combining dental calculus with isotope analysis in the Alps: New evidence from the Roman and medieval cemeteries of Lamon, Italy. Quaternary International. 10.1016/j.quaint.2021.11.022

Food and Agriculture Organization of the United Nations (FAO) (Ed.). (1995). Sorghum and millets in human nutrition. Food and Agriculture Organization of the United Nations.

Formichella, G., Soncin, S., & Cocozza, C. (2023). Isotòpia: A Stable Isotope Database for Classical Antiquity [Computer software]. 10.48493/m0m0-b436

Formichella, G., Soncin, S., Lubritto, C., Tafuri, M. A., Fernandes, R., & Cocozza, C. (2024). Introducing Isotòpia: A stable isotope database for Classical Antiquity. PLOS ONE, 19(6), e0293717. 10.1371/journal.pone.0293717

Fuller, B. T., Richards, M. P., & Mays, S. A. (2003). Stable carbon and nitrogen isotope variations in tooth dentine serial sections from Wharram Percy. Journal of Archaeological Science, 30(12), Article 12. 10.1016/S0305-4403(03)00073-6

Ganzarolli, G., Alexander, M., Chavarria Arnau, A., & Craig, O. E. (2018). Direct evidence from lipid residue analysis for the routine consumption of millet in Early Medieval Italy. Journal of Archaeological Science, 96, 124–130. 10.1016/j.jas.2018.06.007

Gasparri, S. (2016). Italia longobarda. Il regno, i Franchi, il papato. Laterza.

Gnecchi-Ruscone, G. A., Szécsényi-Nagy, A., Koncz, I., Csiky, G., Rácz, Z., Rohrlach, A. B., Brandt, G., Rohland, N., Csáky, V., Cheronet, O., Szeifert, B., Rácz, T.Á., Benedek, A., Bernert, Z., Berta, N., Czifra, S., Dani, J., Farkas, Z., Hága, T., … Krause, J. (2022). Ancient genomes reveal origin and rapid trans-Eurasian migration of 7th century Avar elites. Cell, 185(8), 1402-1413.e21. 10.1016/j.cell.2022.03.007

Goldstein, S., Hixon, S., Scott, E., Wolfhagen, J., Iminjili, V., Janzen, A., Chritz, K., Sawchuck, E., Ndiema, E., Sealy, J. C., Stone, A., Zoeller, G., Phelps, L. N., & Fernandes, R. (2022). Presenting the AfriArch Isotopic Database. Journal of Open Archaeology Data, 10(0), Article 0. 10.5334/joad.94

Gyulai, F. (2014). The history of broomcorn millet (Panicum miliaceum L.) In the Carpathian-basin in the mirror of archaeobotanical remains II. From the roman age until the late medieval age. Columella : Journal of Agricultural and Environmental Sciences, 1. 10.18380/SZIE.COLUM.2014.1.1.39

Hakenbeck, S. E., Evans, J., Chapman, H., & Fóthi, E. (2017). Practising pastoralism in an agricultural environment: An isotopic analysis of the impact of the Hunnic incursions on Pannonian populations. PLOS ONE, 12(3), Article 3. 10.1371/journal.pone.0173079

Heather, P. (1995). The Huns and the End of the Roman Empire in Western Europe. The English Historical Review, CX(435), 4–41. 10.1093/ehr/CX.435.4

Heinrich, F., Hansen, A. M., & Erdkamp, P. (2021). Roman isotopes and economic meaning: Millets, manure, mobility, marine signals, and Malthus. Archaeological and Anthropological Sciences, 13(3), 44. 10.1007/s12520-021-01276-6

Hermes, T. R., Frachetti, M. D., Doumani Dupuy, P. N., Mar’yashev, A., Nebel, A., & Makarewicz, C. A. (2019). Early integration of pastoralism and millet cultivation in Bronze Age Eurasia. Proceedings of the Royal Society B: Biological Sciences, 286(1910), Article 1910. 10.1098/rspb.2019.1273

Iacumin, P., Galli, E., Cavalli, F., & Cecere, L. (2014). C4-consumers in southern europe: The case of Friuli V.G. (NE-Italy) during early and central middle ages. American Journal of Physical Anthropology, 154(4), Article 4. 10.1002/ajpa.22553

Jacobsen, T. C. (2009). The Gothic War: Romes Final Conflict in the West. Westholme Publishing, U.S.

James, T. K., Rahman, A., McGill, C. R., & Trivedi, P. D. (2011). Biology and survival of broom corn millet (Panicum miliaceum) seed. New Zealand Plant Protection, 64, 142–148. 10.30843/nzpp.2011.64.6013

Jarnut, J. (1982). Geschichte der Langobarden. Kohlhammer.

Jørkov, M. L. S., Heinemeier, J., & Lynnerup, N. (2009). The petrous bone-A new sampling site for identifying early dietary patterns in stable isotopic studies. American Journal of Physical Anthropology, 138(2), Article 2. 10.1002/ajpa.20919

Kirleis, W., Dal Corso, M., & Filipović, D. (Eds.). (2022). Millet and What Else?. The Wider Context of the Adoption of Millet Cultivation in Europe. Sidestone Press. https://www.sidestone.com/books/millet-and-what-else

Laffranchi, Z., Mazzucchi, A., Thompson, S., Delgado‐Huertas, A., Granados‐Torres, A., & Milella, M. (2020). Funerary reuse of a Roman amphitheatre: Palaeodietary and osteological study of Early Middle Ages burials (8th and 9th centuries AD) discovered in the Arena of Verona (Northeastern Italy). International Journal of Osteoarchaeology, 30(4), Article 4. 10.1002/oa.2872

Lee-Thorp, J. A. (2008). On Isotopes and Old Bones. Archaeometry, 50(6), Article 6. 10.1111/j.1475-4754.2008.00441.x

Lee-Thorp, J. A., Sealy, J. C., & van der Merwe, N. J. (1989). Stable carbon isotope ratio differences between bone collagen and bone apatite, and their relationship to diet. Journal of Archaeological Science, 16(6), Article 6. 10.1016/0305-4403(89)90024-1

MacKinnon, M. (2019). Consistency and change: Zooarchaeological investigation of Late Antique diets and husbandry techniques in the Mediterranean region. Antiquité Tardive, 27, 135–148. 10.1484/J.AT.5.119548

Mantile, N., Fernandes, R., Lubritto, C., & Cocozza, C. (2023). IsoMedIta: A Stable Isotope Database for Medieval Italy. Research Data Journal for the Humanities and Social Sciences, 1(aop), 1–13. 10.1163/24523666-bja10032

Marinato, M. (2014). Analisi isotopiche e bioarcheologia come fonti per lo studio del popolamento tra tardo antico e alto medioevo in Italia settentrionale. Dati a confronto per le province di Bergamo, Modena e Verona [PhD]. Università degli Studi di Padova.

Marinato, M. (2017). Analisi degli isotopi stabili delle sepolture altomedievali. In A. Chavarría Arnau (Ed.), Ricerche sul Centro episcopale di Padova: Scavi 2011-2012 (pp. 151–154). SAP, Società archeologica.

Marinato, M. (2018). Potenzialità di un approccio multidisciplinare per lo studio del popolamento antico: Il territorio di Bergamo tra tarda antichità e alto medioevo. In C. Giostra (Ed.), Città e campagna: Culture, insediamenti, economia (secc. VI-IX): II Incontro per l’archeologia barbarica, Milano, 15 maggio 2017. SAP, Società archeologica s.r.l.

Marinato, M. (2019). Alimentazione, Salute e Mobilità della Popolazione in Italia Settentrionale tra IV e VIII Secolo. Approcci Bioarcheologici (SAP Società Archeologica).

Maxwell, A. (2019). Exploring Variations in Diet and Migration from Late Antiquity to the Early Medieval Period in the Veneto, Italy: A Biochemical Analysis [PhD]. University of South Florida.

Miller, N. F., Spengler, R. N., & Frachetti, M. (2016). Millet cultivation across Eurasia: Origins, spread, and the influence of seasonal climate. The Holocene, 26(10), 1566–1575. 10.1177/0959683616641742

Mion, L., André, T., Mailloux, A., Sternberg, M., Morales Muniz, A., Rosello-Izquierdo, E., Llorente Rodríguez, L., & Herrscher, E. (2022). Contribution to Mediterranean medieval dietary studies: Stable carbon and nitrogen isotope data of marine and catadromous fish from Provence (9th–14th CE). Data in Brief, 41, 108016. 10.1016/j.dib.2022.108016

Montanari, M. (1988). Alimentazione e cultura nel Medioevo. Laterza.

Motuzaitė Matuzevičiūtė, G., & Laužikas, R. (2023). A Brief History of Broomcorn Millet Cultivation in Lithuania. Agronomy, 13(8), Article 8. 10.3390/agronomy13082171

Murphy, C. (2016). Finding millet in the Roman world. Archaeological and Anthropological Sciences, 8(1), 65–78. 10.1007/s12520-015-0237-4

Nitsch, E. (2012). Stable isotope evidence for diet change in Roman and Medieval Italy: Local, regional and continental perspectives. [PhD Dissertation]. University of Oxford.

Orfanou, E., Zach, B., Rohrlach, A. B., Schneider, F. N., Paust, E., Lucas, M., Hermes, T., Ilgner, J., Scott, E., Ettel, P., Haak, W., Spengler, R., & Roberts, P. (2024). Biomolecular evidence for changing millet reliance in Late Bronze Age central Germany. Scientific Reports, 14(1), 4382. 10.1038/s41598-024-54824-0

Paladin, A., Moghaddam, N., Stawinoga, A. E., Siebke, I., Depellegrin, V., Tecchiati, U., Lösch, S., & Zink, A. (2020). Early medieval Italian Alps: Reconstructing diet and mobility in the valleys. Archaeological and Anthropological Sciences, 12(3), Article 3. 10.1007/s12520-019-00982-6

Pezo-Lanfranco, L., Mut, P., Chávez, J., Fossile, T., Colonese, A. C., & Fernandes, R. (2024). South American Archaeological Isotopic Database, a regional-scale multi-isotope data compendium for research. Scientific Data, 11(1), 336. 10.1038/s41597-024-03148-9

Plecerová, A., Kaupová, S. D., Šmerda, J., Stloukal, M., & Velemínský, P. (2020). Dietary reconstruction of the Moravian Lombard population (Kyjov, 5th–6th centuries AD, Czech Republic) through stable isotope analysis (δ13C, δ15N). Journal of Archaeological Science: Reports, 29, 102062. 10.1016/j.jasrep.2019.102062

Pohl, W. (2008a). Die Langobarden zwischen Elbe und Italien. In M. Hegewisch (Ed.), Die Langobarden: Das Ende der Völkerwanderung. (pp. 23–33). Primus in Herder.

Pohl, W. (2008b). Migration und Ethnogenesen der Langobarden aus Sicht der Schriftquellen. In J. Bemman & M. Schmauder (Eds.), Kulturwandel in Mitteleuropa: Langobarden, Awaren, Slawen (pp. 1–12). Verlag Dr Habelt.

Pohl, W. (2018). The Avars: A Steppe Empire in Central Europe, 567–822. Cornell University Press.

Ricci, P., Mongelli, V., Vitiello, A., Campana, S., Sirignano, C., Rubino, M., Fornaciari, G., & Lubritto, C. (2012). The privileged burial of the Pava Pieve (Siena, 8th Century AD): The privileged burial of the Pava Pieve (Siena, 8th Century AD). Rapid Communications in Mass Spectrometry, 26(20), Article 20. 10.1002/rcm.6302

Riccomi, G., Minozzi, S., Zech, J., Cantini, F., Giuffra, V., & Roberts, P. (2020). Stable isotopic reconstruction of dietary changes across Late Antiquity and the Middle Ages in Tuscany. Journal of Archaeological Science: Reports, 33, 102546. 10.1016/j.jasrep.2020.102546

Saleh, A. S. M., Zhang, Q., Chen, J., & Shen, Q. (2013). Millet Grains: Nutritional Quality, Processing, and Potential Health Benefits. Comprehensive Reviews in Food Science and Food Safety, 12(3), 281–295. 10.1111/1541-4337.12012

Samtani, R., Mishra, S. S., & Neogi, S. B. (2024). Millets: Small grains, big impact in climate action. The Journal of Climate Change and Health, 20, 100345. 10.1016/j.joclim.2024.100345

Šmerda, J. (2016). The new Lombard burial site of Kyjov in Moravia and its position in the development of the 6th c. AD. In H. Geisler (Ed.), Wandel durch Migration? (pp. 167–179). Verlag Dr Faustus.

Soncin, S., Talbot, H. M., Fernandes, R., Harris, A., Tersch M. von, Robson, H. K., Bakker, J. K., Richter, K. K., Alexander, M., Ellis, S., Thompson, G., Amoretti, V., Osanna, M., Caso, M., Sirano, F., Fattore, L., Colonese, A. C., Garnsey, P., Bondioli, L., & Craig, O. E. (2021). High-resolution dietary reconstruction of victims of the 79 CE Vesuvius eruption at Herculaneum by compound-specific isotope analysis. Science Advances. 10.1126/sciadv.abg5791

Spurr, M. S. (1983). The Cultivation of Millet in Roman Italy. Papers of the British School at Rome, 5, 1–15.

Styring, A. K., Ater, M., Hmimsa, Y., Fraser, R., Miller, H., Neef, R., Pearson, J. A., & Bogaard, A. (2016). Disentangling the effect of farming practice from aridity on crop stable isotope values: A present-day model from Morocco and its application to early farming sites in the eastern Mediterranean. The Anthropocene Review, 3(1), Article 1. 10.1177/2053019616630762

Szpak, P. (2014). Complexities of nitrogen isotope biogeochemistry in plant-soil systems: Implications for the study of ancient agricultural and animal management practices. Frontiers in Plant Science, 5. 10.3389/fpls.2014.00288

Taché, K., Jaffe, Y., Craig, O. E., Lucquin, A., Zhou, J., Wang, H., Jiang, S., Standall, E., & Flad, R. K. (2021). What do “barbarians” eat? Integrating ceramic use-wear and residue analysis in the study of food and society at the margins of Bronze Age China. PLOS ONE, 16(4), e0250819. 10.1371/journal.pone.0250819

Tafuri, M. A., Goude, G., & Manzi, G. (2018). Isotopic evidence of diet variation at the transition between classical and post-classical times in Central Italy. Journal of Archaeological Science: Reports, 21, 496–503. 10.1016/j.jasrep.2018.08.034

Temkina, A. (2021). The Early Medieval Transition: Diet Reconstruction, Mobility, and Culture Contact in the Ravenna Countryside, Northern Italy [MA dissertation]. University of South Florida.

Tian, Y., Koncz, I., Defant, S., Giostra, C., Vyas, D. N., Sołtysiak, A., Pejrani Baricco, L., Fetner, R., Posth, C., Brandt, G., Bedini, E., Modi, A., Lari, M., Vai, S., Francalacci, P., Fernandes, R., Steinhof, A., Pohl, W., Caramelli, D., … Veeramah, K. R. (2024). The role of emerging elites in the formation and development of communities after the fall of the Roman Empire. Proceedings of the National Academy of Sciences, 121(36), e2317868121. 10.1073/pnas.2317868121

Vida, T. (2008). Die Langobarden in Pannonien. In M. Hegewisch (Ed.), Die Langobarden: Das Ende der Völkerwanderung. (pp. 73–89). Primus in Herder.

Ward-Perkins, B. (2006). The Fall of Rome: And the End of Civilization. Oxford University Press.

Wickham, C. (2006). Framing the Early Middle Ages: Europe and the Mediterranean, 400-800. Oxford University Press.

Wickham, C. (2010). The inheritance of Rome: A history of Europe from 400 to 1000. Penguin Books. https://www.overdrive.com/search?q=262A6DC8-859C-4440-91BF-9B911D68217C

Wilkin, S., Ventresca Miller, A., Miller, B. K., Spengler, R. N., Taylor, W. T. T., Fernandes, R., Hagan, R. W., Bleasdale, M., Zech, J., Ulziibayar, S., Myagmar, E., Boivin, N., & Roberts, P. (2020). Economic Diversification Supported the Growth of Mongolia’s Nomadic Empires. Scientific Reports, 10(1), Article 1. 10.1038/s41598-020-60194-0

